# PCO371 intracellular agonism at the parathyroid hormone 1 receptor produces pan-activation of signalling partners

**DOI:** 10.64898/2026.05.27.728112

**Authors:** Farhaan Napier-Khwaja, Zamara Mariam, Yasaman Abdolhay, Theo Redfern-Nichols, Graham Ladds, David R. Poyner, Giuseppe Deganutti, Mark Wheatley, Hoor Ayub

## Abstract

Small-molecule agonists of class B1 G-protein-coupled receptors (GPCRs) remain rare because these receptors typically require large extracellular peptide ligands for activation. PCO371 is a notable exception: an intracellular non-peptide agonist, originally developed for osteoporosis treatment, that activates parathyroid hormone 1 receptor (PTH1R) from the cytoplasmic face of the receptor and is considered Gαs-protein biased. In this study, we compared the functional, pharmacological and structural properties of PCO371 with the canonical extracellular peptide agonist PTH 1-34 at the PTH1R to define the molecular mechanisms underlying PCO371’s unusual signalling profile. By manipulating receptor reserve, we demonstrate that PCO371 is a partial agonist relative to PTH 1-34, contrary to previous observations. Across Gαs, Gαi3, Gαq, cAMP, and β-arrestin-2 pathways, operational model analysis showed that PCO371 is non-biased, engaging the same transducers as PTH1-34 but with ∼1000-fold lower potency. Mechanistic modelling using the Cubic Ternary Complex Activation model revealed that reduced intrinsic efficacy is sufficient to reproduce PCO371’s distinct monotonic signalling kinetics, providing a quantitative explanation for its partial agonism. PCO371 binding is β-arrestin-compatible but only drives measurable β-arrestin-2 recruitment when PTH1R is highly expressed. Molecular modelling suggests that PC0371 stable binding needs an intact receptor-G protein complex. Overall, these insights define the mechanistic basis of intracellular agonism at a class B1 GPCR and provide a framework for designing next-generation small-molecule modulators that exploit this emerging pharmacological space.

**GRAPHICAL ABSTRACT:** 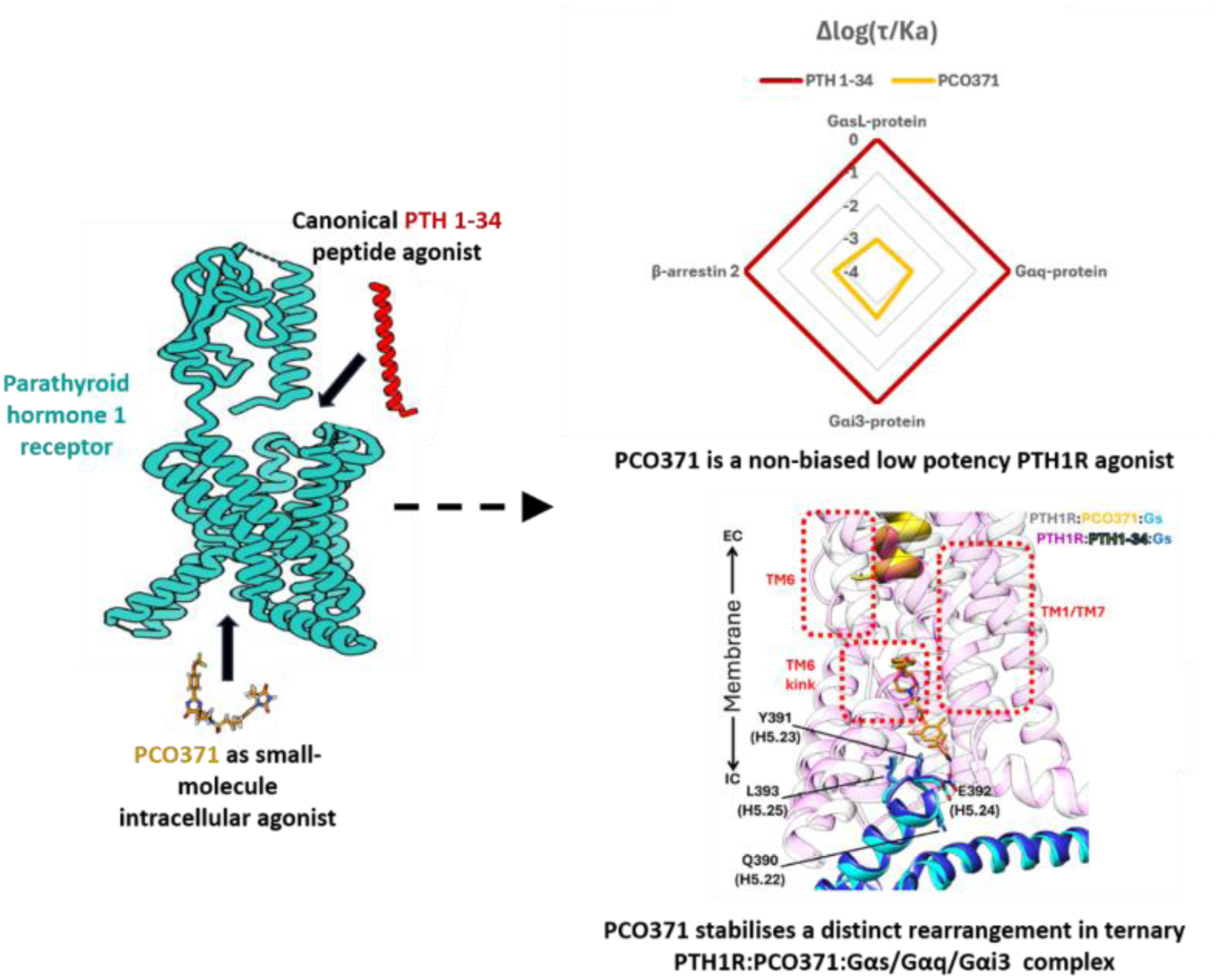

## 1. INTRODUCTION

Class B1 G-protein-coupled receptors (GPCRs) are important therapeutic targets for a wide range of metabolic, hormonal, and age-related disorders (1). These receptors predominantly couple to the stimulatory G-protein (Gαs) to activate adenylyl cyclase but are also capable of activating several distinct signalling pathways (2). Among class B1 GPCRs, the parathyroid hormone type-1 receptor (PTH1R) is a clinically validated target with approved peptide agonists used to treat osteoporosis and hypoparathyroidism, and there is increasing interest in its roles within cardiac and vascular tissues (3–5).

Current therapies rely on peptide agonists such as teriparatide (PTH 1–34). However, these ligands have short plasma half-lives and must be administered by daily subcutaneous injection. Even longer-acting analogues such as eneboparatide still require injection and are rapidly cleared (6). Prolonged receptor activation can also induce desensitisation and lead to side effects, reducing efficacy during chronic treatment. These limitations have driven growing interest in non-peptide agonists for class B1 GPCRs. Small molecules could offer improved stability, oral bioavailability, and more tailored signalling properties, including the potential to minimise desensitisation (7). This has prompted recent efforts to discover and characterise non-peptide agonists with novel mechanisms of receptor engagement (8,9). PCO371 is a non-peptide agonist of the PTH1R developed by Chugai Pharmaceutical (8). Although discontinued in Phase I trial (10), PCO371 remains an important compound for improving small-molecule based receptor agonism. Its discovery emerged from a high-throughput screening platform using LLC-PK1 cells stably expressing approximately one million PTH1R receptors per cell, with a urokinase-type plasminogen activator readout to identify receptor activation (8,11). Unlike peptide ligands, PCO371 does not interact with the receptor’s extracellular N-terminal domain or canonical orthosteric binding pocket. Instead, structural studies demonstrated that it acts as an intracellular “molecular wedge”, interacting with receptor transmembrane (TM) helices, including TM2, TM3, TM6, TM7, Helix 8, and the α5-helix of Gαs-protein (12,13). This unusual binding mode enables activation of Gαs-protein while reportedly avoiding β-arrestin recruitment (8,12). Although it binds at a site distinct from peptide agonists, PCO371 can still antagonise peptide binding (8). Its unique mechanism of activation and proposed Gαs-protein bias make it a compelling candidate for detailed mechanistic investigation.

The aim of this study was to determine how receptor expression level influences PCO371 signalling bias, to provide a quantitative assessment of its efficacy across PTH1R-dependent pathways, and to establish whether PCO371 acts as a biased agonist or a low-efficacy partial agonist. The development of a tetracycline-inducible PTH1R cell line enabled controlled modulation of receptor expression within a single clonal population. This, combined with analysis by the Furchgott method and the Black-Leff Operational Model of Agonism (14,15) provided a robust framework for assessing the contribution of receptor-reserve to signalling (16). Using this analysis we have established whether PCO371’s apparent bias reflects true transducer selectivity or reduced intrinsic efficacy, as observed for other putatively biased ligands such as μ-opioid receptor agonists (17,18). To further support interpretation of ligand-specific signalling behaviour, we also incorporated the Cubic Ternary Complex Activation (CTCA) model (19) as a mechanistic framework for exploring how PCO371-dependent parameters influence G-protein dissociation and reassociation dynamics.

Whilst two structures show how PC0371 binds to the PTH1R (12,13), we have now used molecular-dynamics simulations to further define the activation mechanism of PCO371 at PTH1R. We show that PCO371 requires preassembled receptor-G-protein complexes and high receptor reserve to stabilise a distinct active-state geometry, resulting in full pathway activation at high ligand concentrations but reduced potency compared with PTH 1-34. Activation of Gαi3, and Gαq subtypes was also observed, confirming that PCO371 engages multiple G-protein families without detectable signalling bias.

As PCO371 has been predicted to stimulate several class B1 GPCRs, including wild-type pituitary adenylate cyclase-activating polypeptide type 1 receptor, glucagon receptor, vasoactive intestinal peptide receptor 1 and 2 (12), this work has implications for all of these receptors. Our findings refine current models of class B1 GPCR activation and establish a mechanistic basis for intracellular agonism, providing timely insight into how small molecules can modulate GPCR activity from the cytoplasmic face and informing future design of next-generation therapeutically relevant modulators.

## 2. MATERIALS AND METHODS

### 2.1. Materials

DS69910557 (#HY-148350), PCO371 (#HY-100856), and Parathyroid hormone PTH1-34 amide (#HY-P4821) were purchased from Cambridge Bioscience (Cambridge, UK). Forskolin (FSK) (#1099), and 3-isobutyl-1-methylxanthine (IBMX) (#2845) were purchased from Tocris (Bristol, UK).

### 2.2. Fluorescent PTH 1-34

A custom fluorescent PTH 1-34 analogue was generated by site-specific labelling at Lys¹³, following the strategy previously reported for TAMRA conjugation (20). The modified peptide was conjugated at the ε-amine of Lys¹³ with boron-dipyrromethene (BODIPY) 630/650, yielding a BODIPY-PTH 1-34 ligand. Synthesis was performed by Biosynth using their custom peptide synthesis service. The peptide sequence and characterisation data were previously reported by Khwaja *et al.* (21).

### 2.3. PTH1 receptor cDNA construct design

A NanoLuciferase (NLuc) tagged PTH1R cDNA was generated containing N-terminal His_10_ tag and HA tag followed by the NLuc tag, the human PTH1R(24-593) coding sequence, and a Twin-Strep tag sequence at the 3′ end terminating with a stop codon. The cDNA was custom synthesised by GenScript and cloned into pcDNA 3.1 (+) and pACMV-TetO vector as described by Reeves *et al.* (22). Construct designs and corresponding amino-acid sequences for NLuc-PTH1R were as reported by Khwaja *et al.* (21).

### 2.4. Cell culture and transfection

HEK 293T cells (Public Health England) were cultured in high-glucose Dulbecco’s Modified Eagle Medium (DMEM) supplemented with 10% Fetal Bovine Serum (FBS) and used for experiments between passages 20 and 30. Cells were seeded in T175 flasks and grown to 60-80 % confluence prior to transfection with NLuc-PTH1R pcDNA 3.1 (+) using polyethylenimine (PEI) at a DNA:PEI ratio of 1:3. Transfected cells were cultured for a further 48 h, harvested by centrifugation at 1000 × g, and the resulting pellet stored at –80 °C.

For generation of stable cell line, NLuc-PTH1R cloned in the pACMV-TetO mammalian expression plasmid was transfected into HEK 293S N-acetylglucosaminyltransferase I-deficient (GnTI⁻) tetracycline-repressor (TetR) cells using Lipofectamine 3000. Stable selection was performed by limiting dilution in the presence of geneticin (G418; 2 mg/ml). Monoclonal populations were pharmacologically assessed by cAMP accumulation assays, and functional clones expressing the desired receptor were expanded for downstream characterisation.

HEK 293S GnTI⁻ TetR cells (kind gift from Dr Philip Reeves, University of Essex) stably expressing NLuc-tagged PTH1R or wild-type (WT) PTH1R were seeded in T175 flasks and grown to 60–80 % confluence in DMEM/F12-HEPES supplemented with 10 % FBS and 0.5 mg/ml G418. To induce receptor expression, the medium was replaced with DMEM/F12 containing tetracycline (2 µg/ml) and lacking G418, thereby preventing TetR binding to the TetO operator upstream of the coding sequence. Cells were cultured for a further 48–72 h, harvested by centrifugation at 1000 × g, and the pellets stored at –80 °C.

### 2.5. LANCE Ultra cAMP accumulation

Intracellular cyclic adenosine monophosphate (cAMP) accumulation was measured using the LANCE Ultra cAMP Detection Kit (Revvity), which employs a Time-Resolved Förster Resonance Energy Transfer (TR-FRET)-based homogeneous assay for quantifying cAMP levels.

Tetracycline-inducible HEK293S GnTI⁻_tetR stably transfected cells were seeded (15,000 cells/well) in clear, 96-well plates coated with poly-D-lysine (0.1 mg/ml). 3-5 h after seeding, the medium was replaced with DMEM/F12-HEPES containing tetracycline and incubated at 37 °C, 5 % CO₂ in a humidified incubator for 16-20 h. The medium was aspirated and replaced with Hank’s Balanced Salt Solution (HBSS) supplemented with 0.1 % BSA, 0.5 mM IBMX, and 1 mM HEPES, followed by incubation at 37 °C for 30 min. Serial dilutions of ligand were added and incubated at 37 °C for 30 min. The reaction was terminated by aspirating the stimulation buffer, and cells were placed on ice. For functional-antagonism studies, the agonist was added at a concentration eliciting an EC₈₀ response, followed by serial dilutions of antagonist and incubation for 30 min. The reaction was terminated by aspirating the stimulation buffer, and cells were placed on ice.

50 μl of ice-cold absolute ethanol was added and evaporated off at 37 °C for 2 h. 75 μl of lysis buffer (0.35 % Triton X-100, 50 mM HEPES, 10 mM CaCl_2_; pH 7.4) was added and incubated for 15 min with shaking (250 rpm). 10 μl of cell lysate was transferred to a white, 384-well Optiplate. A cAMP standard curve was generated alongside each experiment, where 5 μl of a serial dilution of a known concentration of cAMP and 5 μl of lysis buffer is added to the plate instead of cell lysate. 5 μl of 4x Europium-labelled-cAMP is added to the lysate/standard followed by 5 μl of 4x ULight™-dye conjugated anti-cAMP antibody. Plates were incubated at room temperature protected from light, with gentle shaking (30 rpm) for 30 min. Fluorescence emission at λ = 665 nm was measured using a CLARIOstar^®^ plus plate reader (BMG LabTech) equipped with an excitation filter (λ = 340-20 nm), a dichroic long-pass filter and two emission filters: λ = 665-10 nm and λ = 620-10 nm.

### 2.6. HTRF β-arrestin 2 recruitment

β-arrestin-2 recruitment was measured using the Human Total β-Arrestin-2 Detection Kit (Revvity) based on the Homogeneous Time-Resolved Fluorescence (HTRF) principle. Cells were seeded (100,000 cells/well) in a white, 96-well optiplate coated with poly-D-lysine (0.1mg/ml). 3-5 h after seeding, the media was replaced with DMEM/F12-HEPES containing tetracycline and incubated at 37 °C, 5 % CO_2_ in a humidified incubator, for 16-20 h. The media was aspirated, replaced with serial dilution of ligand prepared in stimulation buffer supplied with the kit, followed by incubation at 25 °C for 30 min. The reaction was terminated by aspirating the stimulation buffer and adding stabilisation buffer for 15 min at 25 °C. Cells were washed three times with wash buffer 1, then incubated overnight at room temperature with pre-mixed d2-labelled and Europium cryptate-labelled antibodies. Fluorescence emission was recorded at λ = 665 ± 10 nm and λ = 620 ± 10 nm using a CLARIOstar® Plus plate reader. HTRF signals were expressed as TR-FRET ratio units (665/620 × 10,000; arbitrary fluorescence units) according to the manufacturer’s instructions, and Eₘₐₓ values were derived from nonlinear regression of concentration-response curves.

### 2.7. TRUPATH G-protein Bioluminescence Resonance Energy Transfer (BRET) assay

The TRUPATH assay was performed as previously described (23) with minor modifications. Tetracycline-inducible HEK293S GnTI^-^_tetR cells were seeded into 10 cm^2^ dishes (700,000 cells/ml) and incubated at 37 °C, 5 % CO_2_ in a humidified incubator for 5-8 h. Cells were transfected with the Gα-Rluc8, Gβ, and GFP2-Gγ biosensor combinations as described (24) using TransIT^®^-2020 according to manufacturer’s instructions. The transfection mixture was supplemented with tetracycline (2 μg/ml) and applied to the cells, followed by incubated at 37 °C, 5 % CO_2_ for 20-24 h. Transfected and induced cells were then dislodged and seeded (70,000 cells/well) into white, 96-well Optiplates and incubated at 37 °C, 5 % CO_2_ for 16-20 h. The media was aspirated and replaced with HBSS supplemented with 0.1 % BSA and 1 mM HEPES. Coelenterazine-400a (7.5 μM) diluted in assay buffer was added and incubated at room temperature for 5 min. Serial dilution of ligand was added, and luminescence was recorded every minute for 15 min at λ = 410 ± 80 nm and λ = 515 ± 30 nm using a CLARIOstar® Plus plate reader.

### 2.8. NanoBRET equilibrium saturation ligand-binding assays

Saturation ligand-binding assays were performed in white, 384-well OptiPlates in a final volume of 50 µl. Serial dilutions of BODIPY-PTH 1-34 were prepared with or without an excess of unlabelled ligand (100 µM PCO371 or 10 µM DS69910557) to define non-specific binding. The reaction was initiated by adding receptor material (0.02–0.2 nM) and incubated at 25 °C for 90 min to establish equilibrium. Furimazine (10 µM; 5 µl) was then added and incubated at 25 °C for 5 min.

NanoBRET measurements were performed using a CLARIOstar® Plus plate reader equipped with the LVF Monochromator™. Emission profiles were recorded sequentially per well at λ = 460 ± 80 nm for the NLuc donor and λ = 660 ± 100 nm for BODIPY-630/650.

### 2.9. NanoBRET equilibrium competition ligand-binding assays

Competition ligand-binding assays were performed in white, 384-well OptiPlates in a final volume of 50 µl containing BODIPY-tracer ligand, stated concentrations of competing ligand, with or without mini-Gα protein, and receptor material (0.02–0.2 nM). Plates were incubated at 25 °C for 90 min. Non-specific binding was determined using a saturating concentration of unlabelled ligand. Furimazine (10 µM; 5 µl) was added and incubated at 25 °C for 5 min. NanoBRET measurements were performed using a CLARIOstar® Plus plate reader. Emission profiles were recorded sequentially per well at λ = 460 ± 80 nm for the NLuc donor and λ = 660 ± 100 nm for BODIPY-630/650.

### 2.10. PCO371 force field parameterisation

PCO371 was modelled in two different keto-enol tautomeric states keto A and enol tautomers, (Supplementary Figure 6) and submitted to CGenFF (25) to retrieve force filed topologies and parameters. The two tautomers were simulated in complex with PTH1R (without Gαs) as reported below. After equilibration, the tautomer that displayed lower RMSD (keto tautomer) was retained for the successive steps of the study.

### 2.11. PTH1R and G-protein systems modelling

To preserve the precise non-peptide agonist-induced conformation of the receptor, the experimental coordinates of PTH1R from the cryo-EM structure of the PCO371-bound PTH1R-Gαs complex (PDB ID: 8JR9) were utilised.

The nucleotide sequences for the Gα-protein constructs (Gαs, Gαi3, and Gαq) used in the TRUPATH assays were translated into amino acid sequences using the ExPASy Translate tool (https://web.expasy.org/translate/). The translated sequence of the Gαs, Gαi3, and Gαq subunits was modelled separately in complex with the PTH1R using AlphaFold3 (AF3) (26). The AF3-generated Gαs, Gαi3, and Gαq subunits were structurally superimposed onto the experimental Gα coordinates from PDB 8JR9, displaying excellent structural overlap. The alpha-helical domain in the Gα structure was removed from the AF3 model to maintain consistency with the experimental data.

Gβ subunit was derived from PDB 8JR9, while the Gγ subunit was modelled using AF3 to resolve missing regions. The AF3 model was structurally aligned to the N-terminus of the experimental Gγ subunit in PDB 8JR9. The experimental coordinates were retained for the majority of the subunit, while the missing C-terminal residues were sourced from the AF3 prediction. The initial AF3-predicted C-terminal tail exhibited an unrealistically linear conformation and was manually adjusted (bent) to assume a physiologically relevant orientation suitable for membrane anchoring.

The Gα-subunits were palmitoylated at the N-terminal cysteine residue, i.e., Cys2 in Gαs, Cys3 in Gαi3 and Cys9 in Gαq systems. Additionally, a geranylgeranylation modification was introduced to Cys68 of the Gγ subunit.

The final refined components, comprising the PTH1R, small-molecule agonist PCO371, Gβ subunit derived from PDB 8JR9, Gγ and modelled Gs subunits, were assembled in three different systems (i.e., PTH1R:PCO371-Gαs, PTH1R:PCO371:Gαi3 and PTH1R:PCO371:Gαq) using 8JR9 as a template for superimposition.

### 2.12. Molecular dynamics simulations of PTH1R in complex with Gαs, Gαi3 and Gαq

The resulting models of PTH1R:PCO731:Gα_s_, PTH1R:PCO731:Gα_i3_ and PTH1R:PCO731:Gα_q_ were oriented to the membrane via superimposition with the PTH1R receptor in PDB 8FLQ in the OPM^1^database. The complexes were then parameterised with the CHARMM36^2^ force field, and the resulting systems were prepared for simulations via in-house scripts that utilise both Python HTMD^3^ and Tool Command Language (TCL) scripts.

Preliminary hydrogen atoms were added through an extensive multistep procedure employing the pdb2pqr^4^ and PROPKA3^5^ software, considering a simulated pH of 7.4. Following this, the PTH1R:PCO731 complex (i.e. in the absence of any G protein) was embedded in a rectangular 116 Å x 116 Å 1-palmitoyl-2-oleyl-sn-glycerol-3-phosphocholine (POPC) bilayer (previously built by using the VMD Membrane Builder plugin 1.1 at http://www.ks.uiuc.edu/Research/vmd/plugins/membrane/), while the receptors in the Gα_s_, Gα_i3_ and Gα_q_ systems, were embedded in a 146 Å x 146 Å POPC bilayer. The coordinates retrieved from the OPM database were employed in this step, to gain the correct orientation within the membrane, while removing the lipid molecules overlapping the receptor TMD bundle.

TIP3P water molecules^6^ were added to the simulation box; the final dimensions for the four systems were 116 Å x 116 Å x 206 Å (in the absence if any G-protein), and 146 Å x 146 Å x 216 Å (in the presence of G-proteins), using the VMD Solvate plugin 1.5 (VMD Solvate plugin, Version 1.5; http://www.ks.uiuc.edu/Research/vmd/plugins/solvate/). Finally, the overall charge neutrality was reached by adding Na^+^/Cl^−^ counterions (final ionic concentration of 0.150 M) using the VMD Autoionize plugin 1.3 (Autoionize Plugin, Version 1.3; http://www.ks.uiuc.edu/Research/vmd/plugins/autoionize/). For all three systems, the equilibration and production simulations were computed using the ACEMD3^7^ MD engine. Initial steric clashes were mitigated using 2500 steps of conjugate-gradient energy minimisation. Following minimisation, the systems were equilibrated through a series of sequential phases under isothermal-isobaric conditions (NPT), in triplicate (i.e., three separate equilibrations were ran for each system). The temperature (310 K) and pressure (1 atm) during this phase were regulated using a Langevin thermostat^8^ with a damping coefficient of 0.1 ps^-1^ and a Monte Carlo barostat (27), respectively, utilising a 2 fs integration timestep.

In order to resolve clashes, an initial 6 ns MD run was conducted where a harmonic positional restraint of 1 kcal mol^−1^ Å^−2^ was applied to the protein coordinates and the lipid phosphorus headgroups, which was linearly scaled down to zero over the course of the simulation. Subsequent 100 ns of MD simulation were performed, during which restraints were maintained strictly on the protein atoms to allow complete relaxation of the surrounding lipid bilayer and solvent environment. Lastly, positional constraints were applied to the protein backbone Ca atoms and the PCO371 small molecule for a further 20 ns.

A single unrestrained productive replica was subsequently executed for each of the three equilibrations per system, in the canonical ensemble (NVT) for 1 microsecond (for each replica), with an integration time step of 4 fs. The temperature was set at 310 K, using a thermostat damping of 0.1 ps^-1^ and the M-SHAKE algorithm^10^ to constrain the bond lengths involving hydrogen atoms.

For non-bonded interactions, the cut-off distance was set at 9 Å, with a switching function applied beyond 7.5 Å. Long-range Coulomb interactions were handled using the particle mesh Ewald summation method (PME)^11^ by setting the mesh spacing to 1.0 Å.

### 2.13. Molecular dynamics analysis

Root mean square deviations (RMSDs) were calculated across the trajectory frames using Visual Molecular Dynamics (VMD)^12^. Interatomic contacts and hydrogen bonds were detected using the GetContacts scripts tool (https://getcontacts.github.io), setting a hydrogen bond donor-acceptor distance of 3.3 A and a minimum donor-hydrogen-acceptor angle cut-off of 120 degrees. Contacts and hydrogen bond persistences are quantified as the percentage of the MDS frames (over all the frames obtained by merging three different replicas) in which protein residues formed contacts or hydrogen bonds with the ligand.

### 2.14. Mechanistic modelling of PTH1R

To assess the relationship between G-protein reassociation and ligand efficacy, quasi-random sampling was combined with mechanistic modelling. The previously published Cubic Ternary Complex Activation (CTCA) mechanistic model (19,28,29) was chosen as it allows quantification of G-protein dissociation (Biomodel ID: MODEL2306220001) (30). The previously published parameter set for the PTH1R was used to define receptor and G-protein properties in the model, while ligand properties were varied by changing the ligand parameters *k_L_*_+_, *k_L_*_−_, *α*_+_, *α*_−_, *γ*_+_ and *γ*_−_. Ligand properties were quasi-randomly sampled using the Sobol method (31) within bounds defined around the original values as shown in Table 1.

**Table 1:**
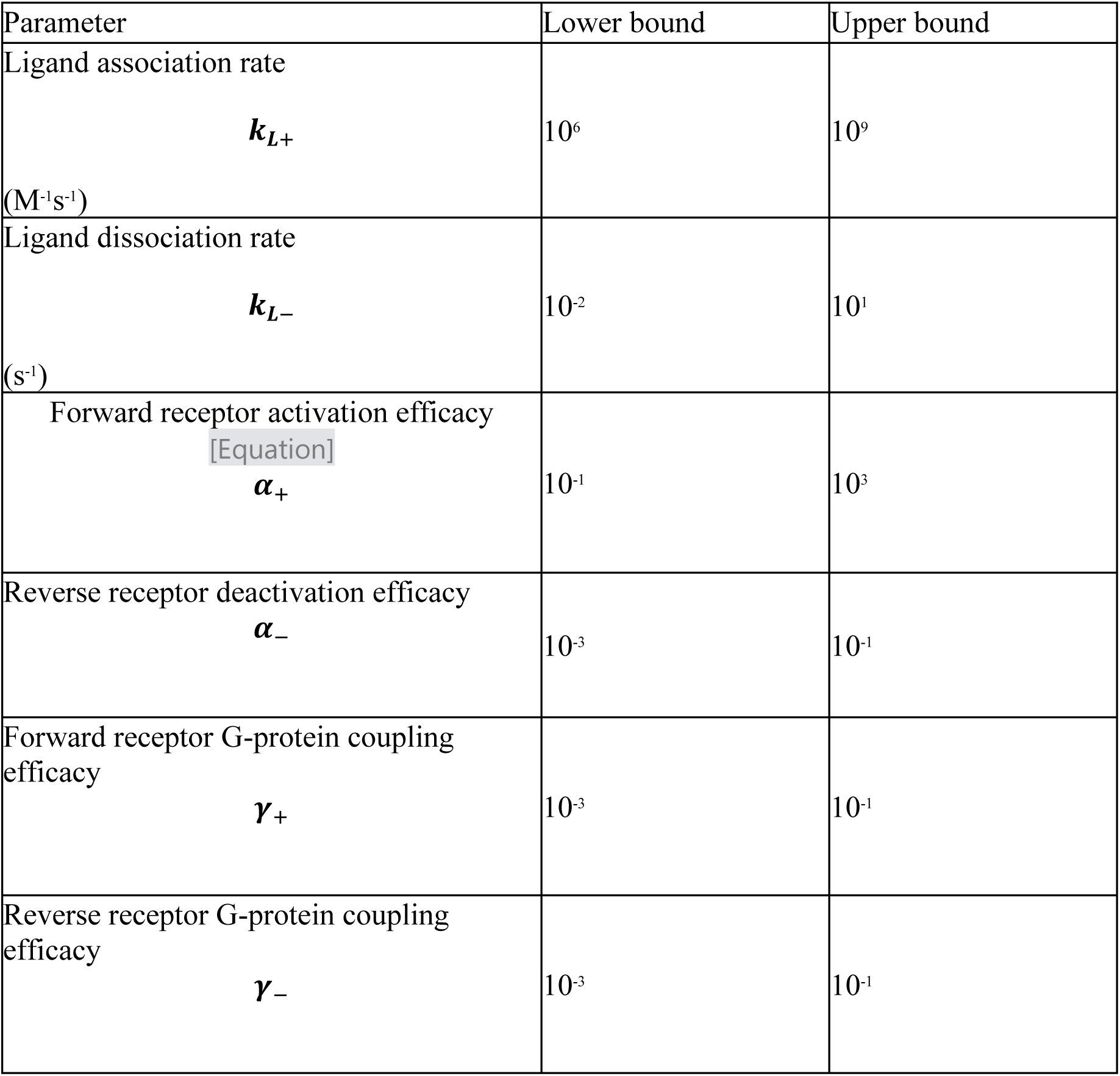
Bounds used for quasi-random sampling of ligand parameters in the Cubic Ternary Complex Activation model. The ligand association (*k*_+_) and dissociation (*k_L_*_−_) rate constants define ligand-receptor binding kinetics. The forward (*α*_+_) and reverse (*α*_−_) receptor activation parameters describe ligand-dependent transitions between inactive and active receptor states. The forward (*γ*_+_) and reverse (*γ*_−_) G-protein coupling parameters describe ligand-dependent G-protein binding to the receptor. These bounds were used to generate quasi-random Sobol samples for mechanistic simulations of ligand behaviour at PTH1R.

| Parameter | Lower bound | Upper bound |
| --- | --- | --- |
| Ligand association rate<br>$k_{L+}$<br>($M^{-1}s^{-1}$) | $10^6$ | $10^9$ |
| Ligand dissociation rate<br>$k_{L-}$<br>($s^{-1}$) | $10^{-2}$ | $10^1$ |
| Forward receptor activation efficacy<br>[Equation]<br>$\alpha_+$ | $10^{-1}$ | $10^3$ |
| Reverse receptor deactivation efficacy<br>$\alpha_-$ | $10^{-3}$ | $10^{-1}$ |
| Forward receptor G-protein coupling efficacy<br>$\gamma_+$ | $10^{-3}$ | $10^{-1}$ |
| Reverse receptor G-protein coupling efficacy<br>$\gamma_-$ | $10^{-3}$ | $10^{-1}$ |

2,048 parameter sets with varying ligand properties were generated in Python (version 3.12.11) using the Quasi-Monte Carlo module of SciPy (version 1.17.1) (32) set to the Sobol method (31). These parameter sets were simulated using COPASI (version 4.46 (build 300)) (33) via its Python interface, BasiCO (version 4.46.300) (34).

Simulations were run for 10^7^ seconds before addition of ligand to allow for equilibration of receptor and G-protein. G-protein dissociation was measured as the sum of *Gα_GTP_* and *Gα_GDP_*. The difference between the maximum G-protein dissociation reached and G-protein dissociation after 30 min of stimulation with 1 µM ligand was calculated to quantify reassociation of G-protein for each parameter set. Representative kinetic traces for two example ligands were generated by selecting individual parameter sets and simulating across further concentrations.

### 2.15. Pharmacological data analysis and statistics

cAMP, β-arrestin-2 and TRUPATH concentration-response curves were fitted using nonlinear regression (four-parameter logistic model) to derive pEC₅₀ and Eₘₐₓ values. TRUPATH kinetic traces were integrated to obtain negative area-under-the-curve (-AUC) values for G-protein activation. NanoBRET competition binding data were analysed using one-site competitive models to determine pKᵢ values under uncoupled and G-protein-coupled conditions. Functional affinity (pKₐ) was calculated using the Operational Model of agonism and the Furchgott–Burzstyn equi-effective concentration method under receptor-reserve modulation.

All data are expressed as mean ± SEM from ≥3 independent experiments performed in three technical replicates. Statistical significance was assessed using one-way ANOVA with Dunnett’s post hoc test (p < 0.05). Curve fitting and statistical analyses were performed using GraphPad Prism (v10.0.3).

## 3. RESULTS

### 3.1. PCO371 activates PTH1R via an antagonist-resistant conformation lacking the high-affinity ternary complex

We investigated whether the antagonist DS69910557 (35) differentially inhibited PTH1R signalling generated by the peptide agonist (PTH 1-34) versus the small-molecule non-peptide agonist (PCO371). PTH1R-expressing cells were stimulated with PTH 1-34 or PCO371 at their respective EC₈₀ concentrations (0.3 nM and 1 µM, respectively) and concentration-inhibition cAMP curves for DS69910557 were generated (Figure 1A-B).

**Figure 1.**
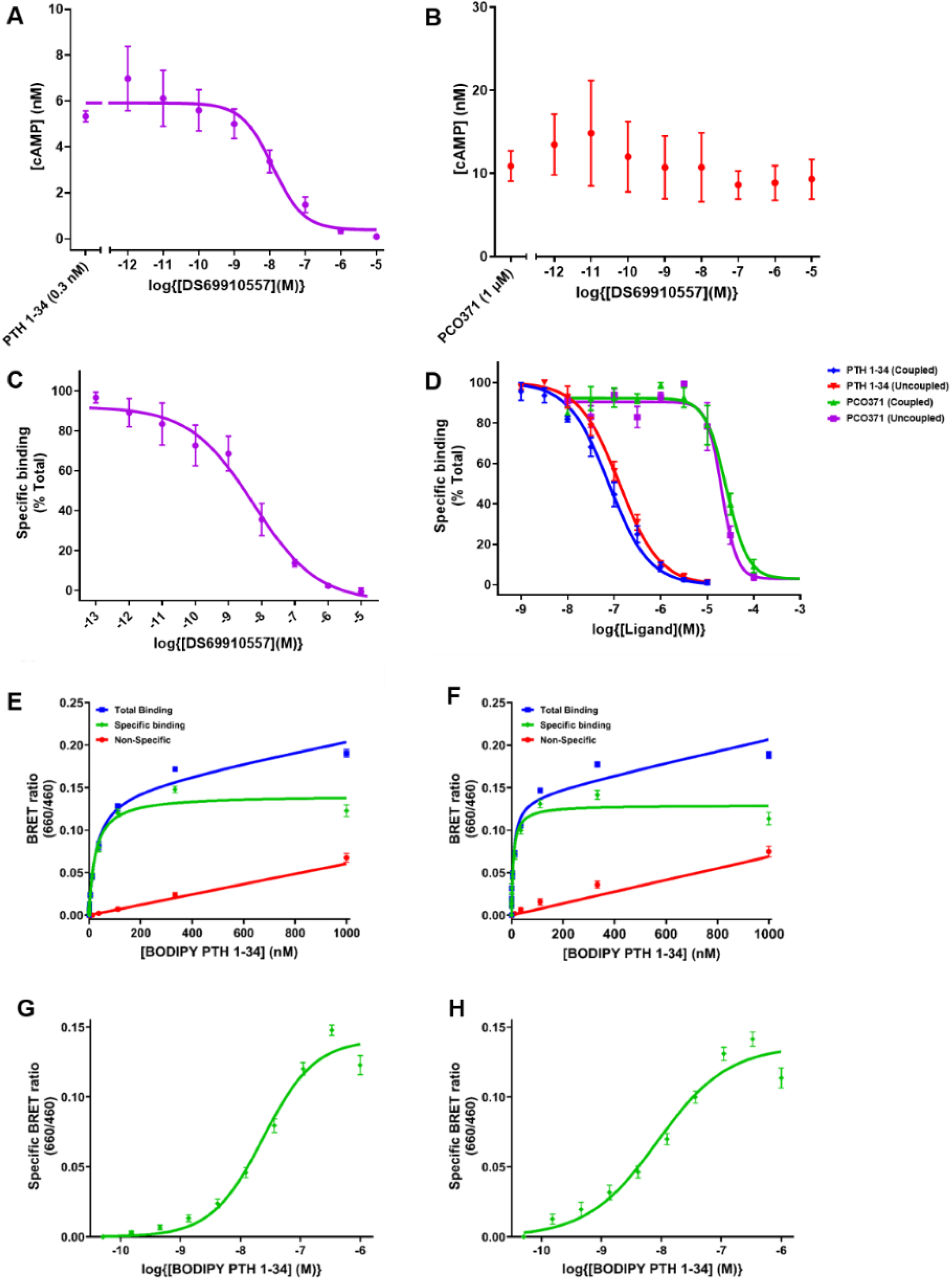
DS69910557 antagonist selectivity confirms distinct receptor engagement by PCO371 for PTH1R. **A)** Concentration–inhibition curve for DS69910557 antagonism of PTH1-34-stimulated cAMP production. PTH1R-expressing cells were stimulated with PTH1-34 (0.3 nM; EC₈₀) and incubated with increasing concentrations of DS69910557 (1 pM–10 µM). **B)** Concentration–inhibition curve for DS69910557 antagonism of PCO371-stimulated cAMP production. PTH1R-expressing cells were stimulated with PCO371 (1 µM; EC₈₀) and incubated with increasing concentrations of DS69910557 (1 pM–10 µM). **C)** Nano-BRET competition binding showing displacement of BODIPY-PTH 1-34 (10 nM) by DS69910557 using NLuc-PTH1R expressing membranes, confirming high-affinity orthosteric engagement. **D)** Nano-BRET competition binding of PTH 1-34 and PCO371 in uncoupled (+Gpp(NH)p, 100 µM) and G-protein-coupled (+mini-Gαs, 3 µM) receptor states. PTH 1-34 displayed a G-protein-dependent increase in affinity, whereas PCO371 affinity was independent of G-protein coupling. Data are mean ± SEM, n = 3, each performed in technical triplicate. (**E-H**) Nano-BRET equilibrium saturation binding shows specific binding of BODIPY-PTH 1-34 to both uncoupled (R₀) and G-protein-coupled (Rᴳ) NLuc-PTH1R conformations. Total binding (n), Specific binding (u), and non-specific binding (λ) defined by DS69910557 (10 μM). **E**) Saturation binding of BODIPY-PTH1-34 in the presence of the non-hydrolysable GTP analogue Gpp(NH)p (100 µM), representing the uncoupled R₀ state. **F**) Saturation binding of BODIPY-PTH1-34 in the presence of mini-Gαs (3 µM), representing the G-protein-coupled Rᴳ state. (**G–H**) Specific binding plotted as log-sigmoidal binding isotherms for R₀ and Rᴳ, respectively. Data are mean ± SEM, n = 3 (technical triplicate). The Rᴳ state displayed significantly higher affinity than R₀ (R₀ pKd = 7.62 ± 0.08; Rᴳ pKd = 8.13 ± 0.11; p = 0.02, unpaired t-test with Welch’s correction).

DS69910557 potently antagonised PTH 1-34-stimulated cAMP response (pIC₅₀ = 7.91 ± 0.03) but in contrast, DS69910557 did not inhibit PCO371-mediated signalling (Figure 2B). This selective inhibition of PTH 1-34 signalling over PCO371 signalling exhibited by DS69910557 is entirely consistent with PCO371 activating PTH1R through a distinct intracellular mode of receptor engagement to PTH 1-34.

**Figure 2.**
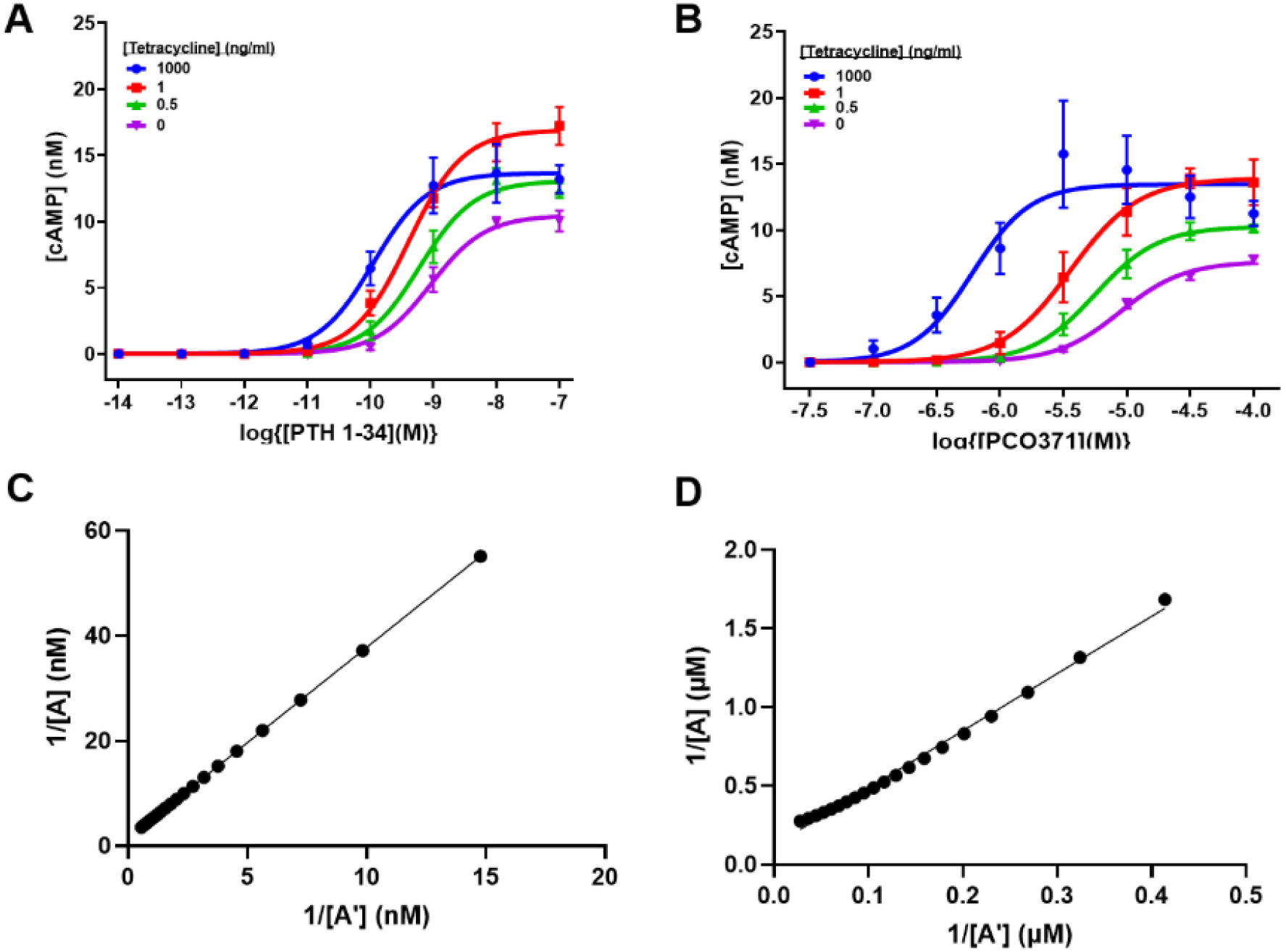
Modulation of PTH1R receptor reserve shows PCO371 dependence on high receptor reserve for full agonism. cAMP accumulation measured at receptor reserves induced by tetracycline concentrations 1000 ng/ml (λ), 1 ((ν), 0.5 (p), and 0 (q) ng/ml. Tetracycline concentrations were empirically optimised to span the full induction range of tetracycline-regulated PTH1R expression, with 0 and 0.5 ng/ml producing very low receptor expression, 1 ng/ml marking minimal induction, and 1000 ng/ml achieving maximal expression. **A**) Concentration-response curves for PTH1-34-stimulated cAMP accumulation. **B**) Concentration-response curves for PCO371-stimulated cAMP accumulation. Data are mean ± SEM, n = 3, performed in technical triplicate. Responses obtained under the 1 ng/ml and 0 ng/ml tetracycline conditions (low-receptor reserve range) were used to derive functional affinity values by plotting the reciprocal of the equi-effective concentrations 1/[A] at 1 ng/ml and 1/[A’] at 0 ng/ml. Functional affinity was also estimated by fitting the Operational Model with receptor depletion. **C)** Functional affinity analysis for PTH 1-34. **D)** Functional affinity analysis for PCO371.

Nano-BRET competition ligand binding assays were performed to test whether DS69910557 engaged the orthosteric peptide-binding pocket of the PTH1R (Figure 1C). Using BODIPY-PTH 1-34 (10 nM) as the fluorescent tracer, DS69910557 displaced the tracer with high affinity (pKi = 8.33 ± 0.11; n= 3), confirming selective competition at the extracellular orthosteric site. To compare this engagement with PTH 1-34 and PCO371, competition binding assays were performed under uncoupled (R₀) and G-protein-coupled (Rᴳ) conditions (Figure 1D; equilibrium saturation binding shown in Figure 1E-H). PTH 1-34 displayed a clear G-protein-dependent affinity shift (R₀ pKi = 7.15 ± 0.02; Rᴳ pKi = 7.52 ± 0.05; n = 3; p = 0.01), consistent with formation of a high-affinity ternary complex upon G-protein coupling. In contrast, PCO371 affinity was independent of the G-protein coupling (R₀ pKi = 4.82 ± 0.06; Rᴳ pKi = 4.94 ± 0.07) with no significant R₀→Rᴳ shift (p = 0.26).

### 3.2. Functional characterisation of PCO371 versus PTH 1-34 at PTH1R

#### 3.2.1 PCO371 displays classical partial-agonist behaviour

PTH1R expression in our engineered cell-line was under tetracycline-inducible regulation, allowing controlled modulation of receptor availability. A series of tetracycline concentrations (0, 0.5, 1, 1000 ng/ml) was used to vary PTH1R expression and thereby generating a graded range of receptor reserve. Intracellular cAMP signalling stimulated by either PTH 1-34 or PCO371 was characterised using cells expressing different PTH1R levels (Figure 2A-B; Table 2). Across these conditions, PTH 1-34 maintained high potency and produced near-maximal cAMP responses even when receptor expression was reduced. In contrast, PCO371 showed a pronounced dependence on receptor reserve, with potency decreasing progressively as receptor expression decreased, and maximal cAMP responses (E_max_) substantially reduced at low receptor expression. Only at the high receptor expression levels, was the Emax for PCO371 comparable to the E_max_ for PTH 1-34, which is consistent with PCO371 being a partial agonist.

**Table 2.** Half-maximal effective concentration (pEC₅₀), maximal response (Eₘₐₓ), and functional affinity (pKₐ) values for PTH 1-34 and PCO371. Values were determined under receptor-reserve modulation to assess their impact on cAMP accumulation. Data represent mean ± SEM (n=3) from independent experiments fitted using the Operational Model of agonism and the Furchgott–Burzstyn analysis.

|  | [Tetracycline] (ng/ml) |  |  |  |  |  |  |  |  |  |
| --- | --- | --- | --- | --- | --- | --- | --- | --- | --- | --- |
|  | 1000 |  | 1 |  | 0.5 |  | 0 |  | Functional affinity (pK<sub>a</sub>) |  |
| Ligand | pEC<sub>50</sub> | E<sub>max</sub> (nM) | pEC<sub>50</sub> | E<sub>max</sub> (nM) | pEC<sub>50</sub> | E<sub>max</sub> (nM) | pEC<sub>50</sub> | E<sub>max</sub> (nM) | Operational model | Furchgott and Burzstyn |
| PTH 1-34 | 9.97 ± 0.15 | 14.01 ± 1.85 | 9.38 ± 0.23 | 18.57 ± 2.44 | 9.23 ± 0.15 | 13.41 ± 0.84 | 9.06 ± 0.17 | 10.85 ± 0.72 | 8.83 | 8.76 |
| PCO371 | 6.19 ± 0.09 | 13.56 ± 2.05 | 5.43 ± 0.11 | 13.93 ± 1.19 | 5.23 ± 0.10 | 10.33 ± 0.37 | 5.01 ± 0.11 | 7.89 ± 0.56 | 5.26 | 4.66 |

Functional affinity estimates from Operational Model of Agonism fits (15) obtained under low-receptor-expression conditions further distinguished the two ligands (Figure 2C-D). PTH 1-34 displayed high functional affinity (pKₐ = 8.83 ± 0.13; n = 3), whereas PCO371 showed markedly lower affinity (pKₐ = 5.26 ± 0.23; n = 3).

#### 3.2.2. PCO371 exhibits detectable β-arrestin-2 recruitment

PCO371 displayed a distinct β-arrestin-2 recruitment profile when assessed under high and low PTH1R expression (Figure 3A). Under high receptor expression, PCO371 produced a measurable β-arrestin-2 response (pEC₅₀ = 4.83 ± 0.05; Eₘₐₓ = 2996 ± 457.4; n = 3; E_max_ values expressed in TR-FRET ratio units (665/620 × 10,000; arbitrary fluorescence units)), indicating that β-arrestin-2 engagement is possible. However, β-arrestin-2 recruitment was not detectable when receptor expression was reduced. In contrast, PTH 1-34 recruited β-arrestin-2 robustly across receptor densities (Figure 3B-C). At high expression, PTH 1-34 displayed a pEC₅₀ of 8.20 ± 0.07 and an Eₘₐₓ of 6382.00 ± 439.89 (n = 3). When receptor expression was reduced, potency remained high (pEC₅₀ = 8.40 ± 0.17), although the maximal response decreased (Eₘₐₓ = 1122.33 ± 56.23; n = 3), consistent with reduced receptor availability. This indicates that β-arrestin-2 engagement by PCO371 is weak but present, and dependent on receptor reserve.

**Figure 3.**
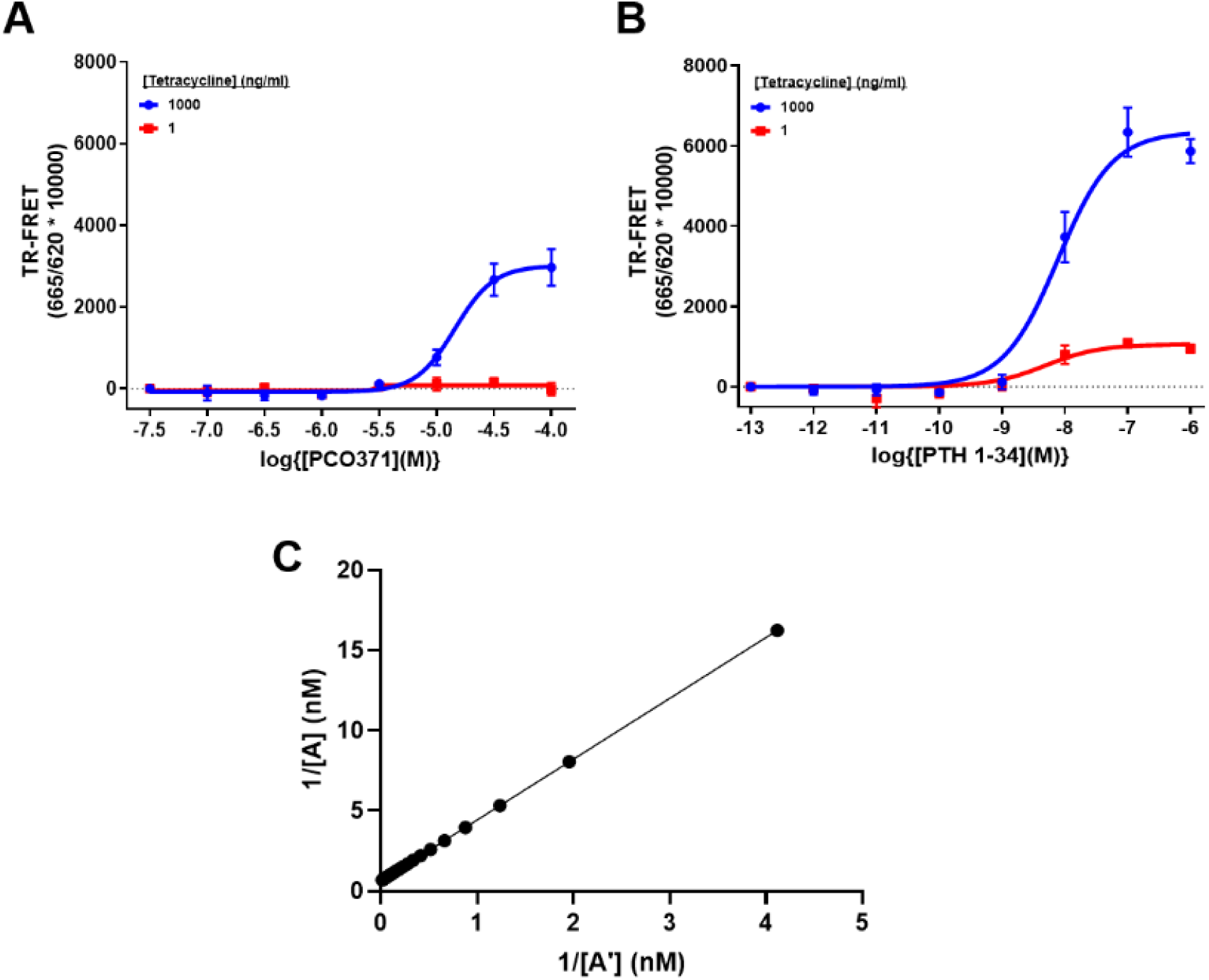
PCO371 functions as a partial agonist for β-arrestin 2 recruitment. Dose-response curves for **A)** PCO371-stimulated or **B)** PTH 1-34-stimulated β-arrestin-2 recruitment using TR-FRET assay at receptor reserves induced by 1000 (λ) or 1 (ν) ng/ml tetracycline. Data are mean ± SEM (n = 3), each performed in technical triplicate. For PTH 1-34, responses obtained under high and low receptor expression conditions were used to derive functional affinity values by plotting the reciprocal of the equi-effective concentrations (1/[A] and 1/[A′]) and by fitting the Operational Model with receptor depletion. **C)** Functional affinity analysis for PTH 1-34, showing the equi-effective concentration plot and corresponding Operational Model fit used to derive pK_a._

#### 3.2.3. PCO371 activates multiple Gα-protein subtypes but with lower potency than PTH 1-34

G-protein activation was assessed using TRUPATH biosensors for GαsL, Gαi3 and Gαq proteins (Figure 4). Both PTH 1-34 and PCO371 promote Gα dissociation from Gβγ for all three subtypes, confirming that PCO371 engages multiple G-protein families rather than signalling selectively through Gαs-protein pathway. WT Gαq-protein produced no measurable TRUPATH signal and was therefore replaced with the stabilised Gαq(R183Q) variant (36). Concentration-response curves derived from negative area under the curve (-AUC) values confirm activation of GαsL, Gαq(R183Q) and Gαi3 by both ligands (Figure 5A-C). Across subtypes, PCO371 is ∼1000-fold less potent than PTH 1 34 (ΔpEC₅₀ ≈ 3; Table 3). Despite this potency difference, maximal responses were comparable at saturating concentrations. This pattern indicates that PCO371 has markedly reduced signalling efficiency, and that its apparent full agonism reflects receptor reserve rather than intrinsic efficacy.

**Figure 4.**
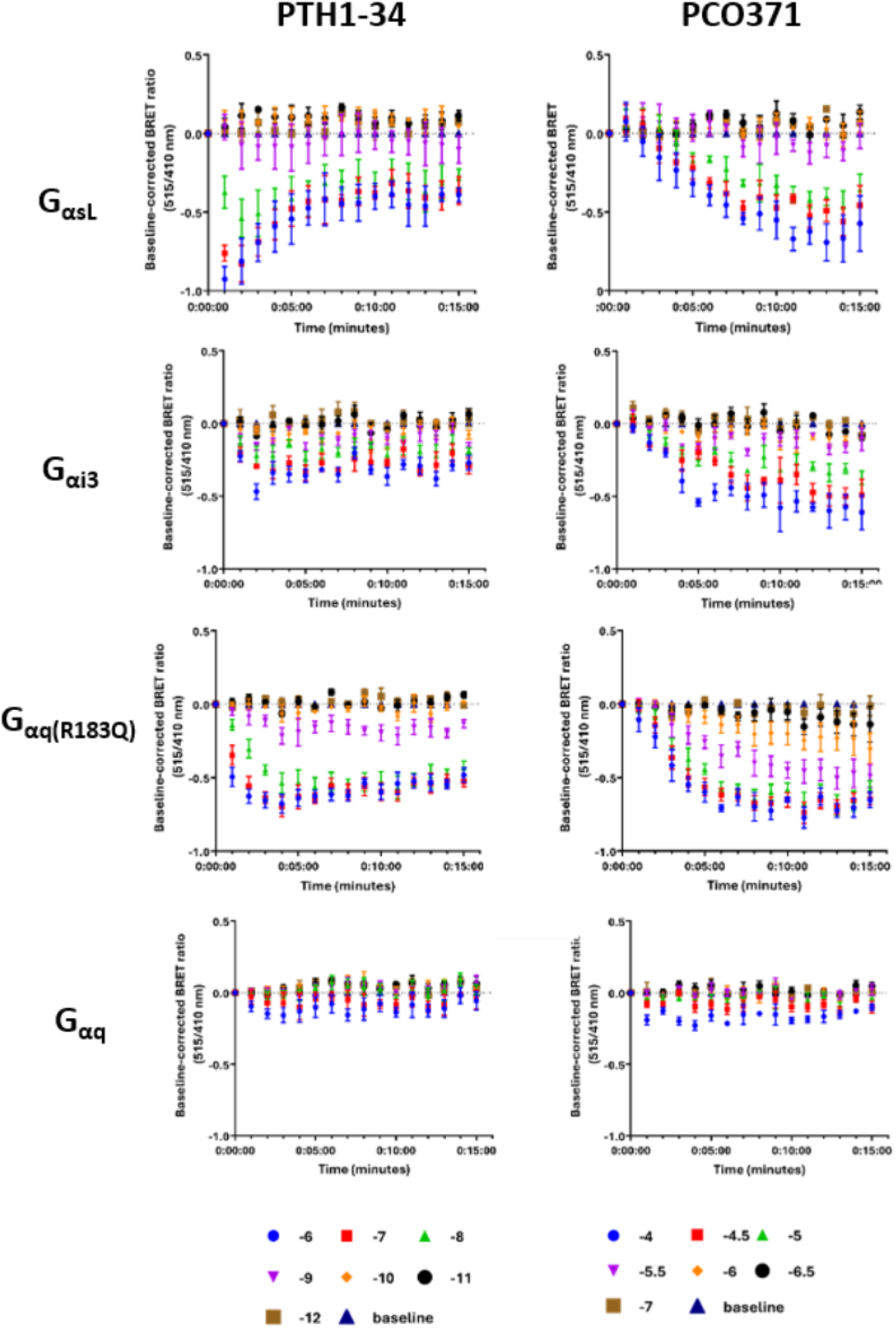
Raw TRUPATH data for PTH1-34 and PCO371 signalling at different Gα-subunits. Tetracycline induced (2 μg/ml) HEK293S GnTI-_tetR cells stably expressing PTH1R cells were transfected with the indicated TRUPATH construct and then stimulated with concentrations shown of PTH 1-34 or PCO371. Data are mean ± SEM, n =3, performed in technical duplicate.

**Figure 5.**
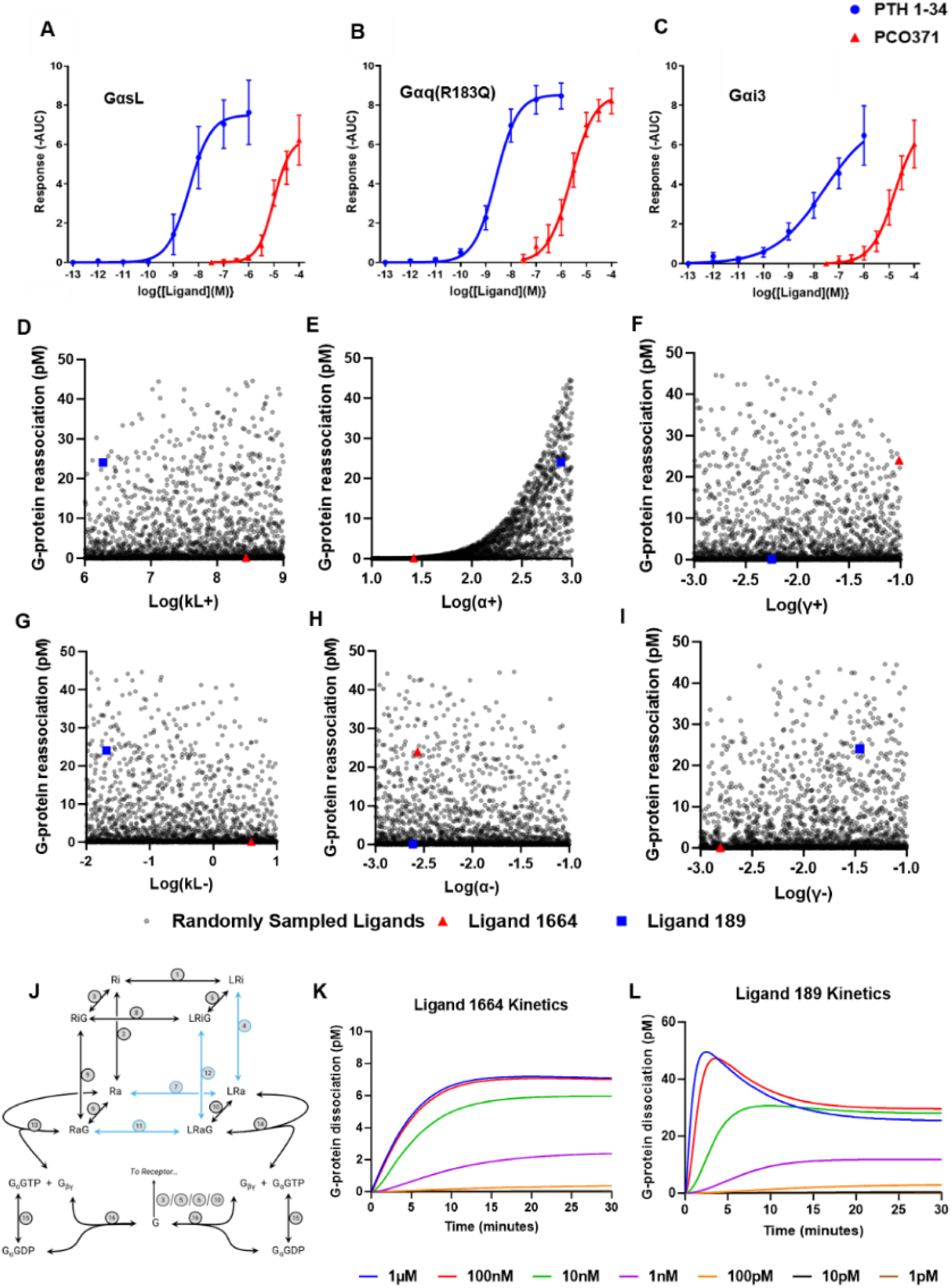
TRUPATH activation shows all Gα-protein subtype engagement by PCO371 and CTCA modelling identifies intrinsic efficacy as sufficient to generate peak-and-fall kinetics. **A-C)** Concentration-response curves (-AUC values) for PTH 1-34 and PCO371 measured using TRUPATH G-protein biosensors in HEK 293S cells stably expressing PTH1R. PTH 1-34-stimulated (λ) or PCO371-stimulated (p) responses are shown for **A)** GαsL **B)** Gαq(R183Q) **C)** Gαi3. Data represent mean ± SEM, n = 3, independent experiments, each performed in technical duplicate. **D-I)** Cubic Ternary Complex Activation (CTCA) modelling output showing quasi-random ligand sampling plotted as G-protein reassociation against logarithmically distributed ligand parameters: **D)** k₊ (ligand association rate), **E)** α₊ (forward receptor-activation efficacy), **F)** γ₊ (forward receptor–G-protein coupling efficacy), **G)** k₋ (ligand dissociation rate), **H)** α₋ (reverse receptor-activation efficacy), and **I)** γ₋ (reverse receptor–G-protein coupling efficacy). **J)** Cubic Ternary Complex Activation Model illustrating all receptor, ligand, and G-protein states and transitions. The cube depicts the eight receptor configurations (Ri, LRi, RiG, LRiG, Ra, LRa, RaG, LRaG) connected by numbered reactions (1–12) representing ligand binding/unbinding, receptor activation/deactivation, and G-protein coupling/uncoupling. Additional reactions (13–16) describe G-protein activation, GαGTP hydrolysis and G-protein reassociation. α-containing reactions highlighted (blue arrows). **(K–L)** Simulated kinetic traces for representative ligands across a range of ligand concentrations (1 pM–1 µM). The PCO371-like kinetics of ligand 1664 in **K)** displayed a slow monotonic rise toward steady state with no peak, whereas the PTH 1-34-like kinetics of ligand 189 in **L)** produced a transient peak followed by partial recovery. Simulations were performed under conditions previously described (19).

**Table 3.** Summary of potency and maximal response values (pEC₅₀ and Emax) for each ligand across the three G-protein subtypes. Data are mean ± SEM, n = 3 performed in technical duplicate using TRUPATH Gα subtype biosensors. Emax values were derived from - AUC measurements of TRUPATH BRET responses.

|  | **GasL** |  | **Gαq(R183Q)** |  | **Gαi3** |  |
| --- | --- | --- | --- | --- | --- | --- |
| **Ligand** | **pEC<sub>50</sub>** | **E<sub>max</sub>**<br>**(-AUC)** | **pEC<sub>50</sub>** | **E<sub>max</sub>**<br>**(-AUC)** | **pEC<sub>50</sub>** | **E<sub>max</sub>**<br>**(-AUC)** |
| **PTH 1-34** | 8.33 ± 0.21 | 7.54 ± 1.37 | 8.58 ± 0.10 | 8.54 ± 0.66 | 7.84 ± 0.14 | 6.48 ± 1.50 |
| **PCO371** | 4.96 ± 0.12 | 6.16 ± 1.43 | 5.62 ± 0.10 | 8.47 ± 0.84 | 4.73 ± 0.12 | 6.04 ± 1.20 |

The kinetic profile differed between the two ligands, particularly for GαsL (Figure 4). PTH 1-34 produced a characteristic peak-and-fall response, also reported previously (19), reflecting rapid heterotrimer dissociation followed by partial reassociation. In contrast, PCO371 generated a slower monotonic rise toward steady state, lacking the transient peak observed for PTH1-34. This distinct kinetic signature motivated Cubic Ternary Complex Activation (CTCA) modelling (19) to identify which receptor-ligand parameters could account for the divergence. CTCA modelling using quasi-random ligand sampling showed that five of the six simulated parameters (Figure 5D-I), *k_L_*_+_, *k_L_*_−_, *γ*_+_, *γ*_−_, and *α*_−_ did not systematically predict peak-and-fall behaviour. In contrast, *α*_+_ (forward receptor activation efficacy) displays a clear trend, with higher *α*_+_ values associated with greater G-protein reassociation. A schematic of the CTCA reaction network, with *α*_+_-dependent transitions highlighted, is shown in Figure 5J. Time-course simulations (Figure 5K-L) confirmed these relationships: the PTH 1-34-like ligand produced a transient peak followed by partial recovery, whereas the PCO371-like ligand showed a slow monotonic rise toward steady state. These modelling results demonstrate that reduced intrinsic efficacy, captured by *α*_+_, is sufficient to reproduce the kinetic divergence observed experimentally. This supports the conclusion that PCO371 behaves as a low-efficacy or partial agonist at PTH1R.

### 3.3 PCO371 is a non-biased agonist across G-protein subtypes and β-arrestin pathways

Because PCO371 has been reported to display signalling bias at the PTH1R (12,13), we applied the classical pharmacological Operational Model of agonism (14,15) to quantify pathway preference. Transduction coefficient Δlog(τ/Kₐ) were generated for each readout (TRUPATH, cAMP, β-arrestin-2) normalised to PTH 1-34 (shown as radar plot in Figure 6A). Using GαsL as the reference pathway, pathway bias values (ΔΔlog(τ/Kₐ)) were then calculated for direct pathway comparisons (Figure 6B). Across all pathways examined, ΔΔlog(τ/Kₐ) values clustered near zero, indicating no consistent pathway preference for PCO371. This contrasts with earlier reports suggesting Gαs-selective signalling (8) and instead proposes a model in which PCO371 engages all tested transducers with uniformly reduced potency relative to PTH 1-34, without significant pathway bias.

**Figure 6:**
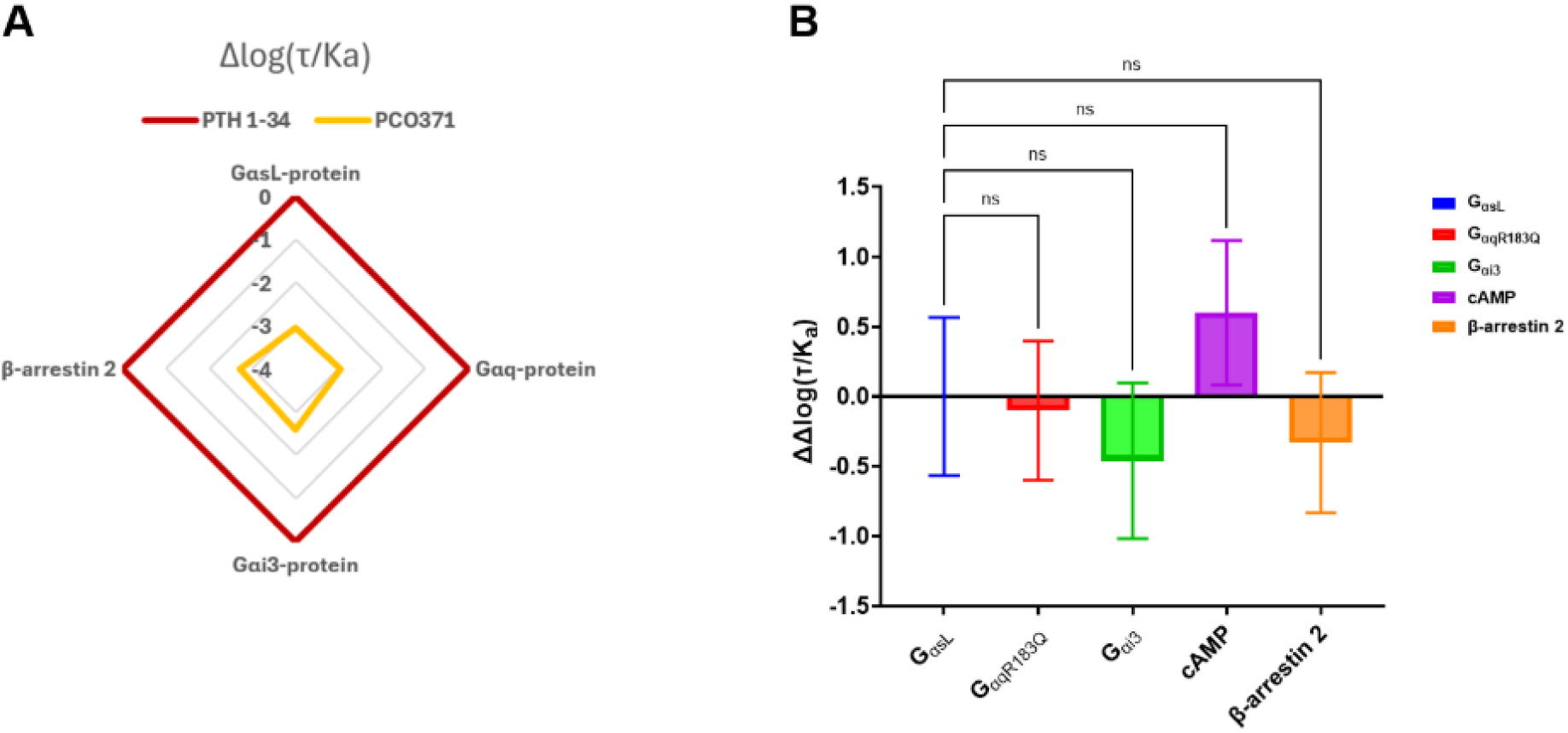
Quantification of PCO371-mediated signalling bias shows no detectable pathway preference at PTH1R. **A)** Radar plot showing transduction coefficients (Δlog(τ/KA)) for PCO371 and PTH 1-34 across GαsL, Gαq, Gαi3 and β-arrestin-2 pathways. **B)** Pathway bias values ΔΔlog(τ/Ka) for PCO371 were calculated using GαsL as the reference pathway and PTH 1-34 as the normalising agonist. Statistical significance was assessed using one-way ANOVA with Dunnett’s post hoc test (p < 0.05)

### 3.4. Structural mechanism of PCO371 action

#### 3.4.1. PCO371 induces a distinct active-state geometry of PTH1R

Comparative structural analysis of PTH1R:Gαs complexes bound to PTH 1-34 or PCO371 reveals pronounced differences in TM6 and adjacent regions (Figure 7A). In the PTH 1-34 complex, TM6 exhibits a sharp kink typical of the canonical active state; in the PCO371 complex, TM6 adopts a straighter, more linear conformation with a short linker that accommodates the small molecule in a boomerang-like conformation. The extracellular region of TM6 becomes fully helical in the PCO371 state, and the TM1–TM7 bundle is less bent than in the PTH 1-34-bound structure. These rearrangements define a distinct active-state geometry stabilised by PCO371 compared to PTH 1-34.

**Figure 7.**
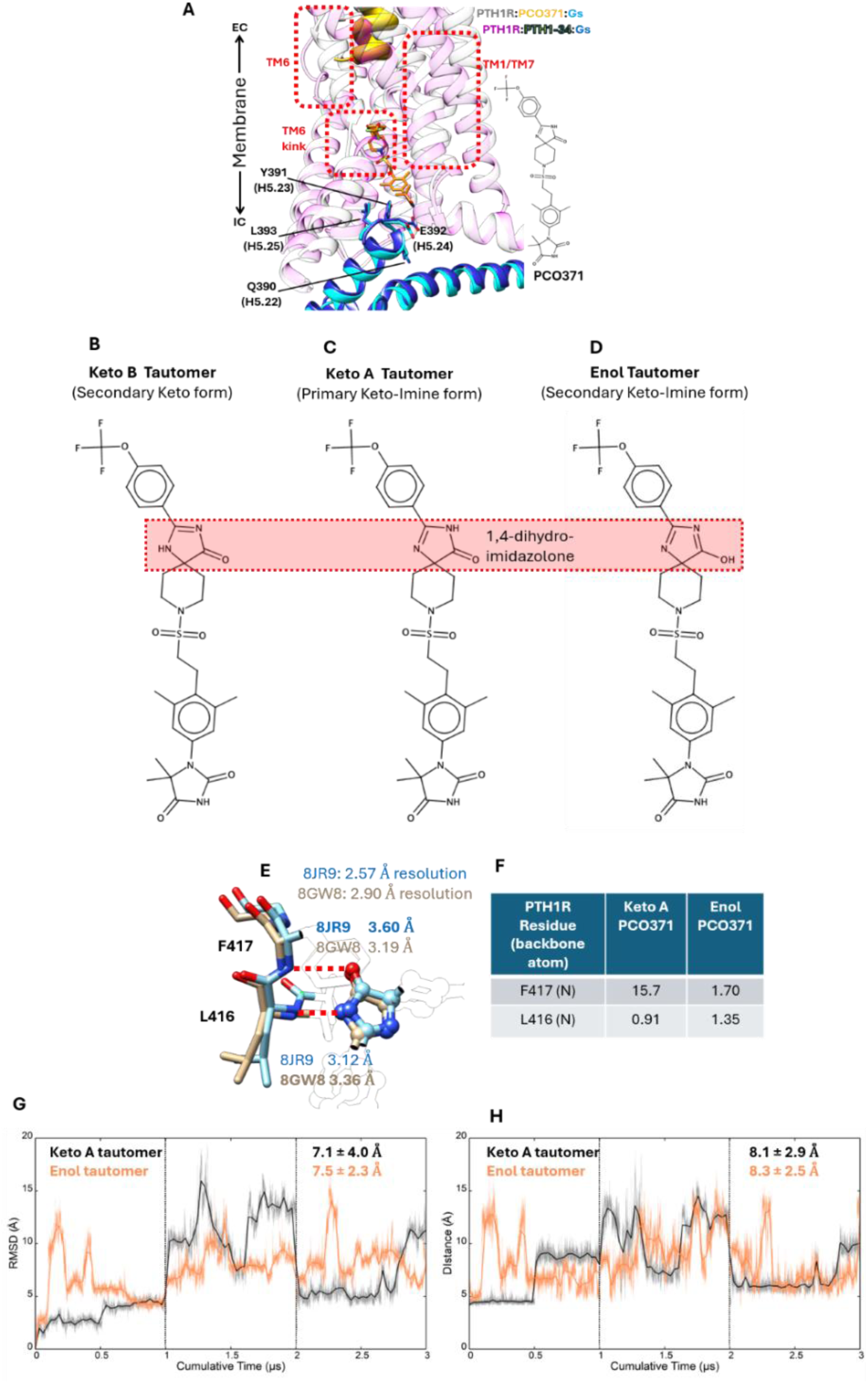
Tautomeric and structural dynamics of PCO371 at the PTH1R binding site. **A)** Comparison between PTH1R in complex with PCO371 (white ribbon, PDB 8JR9) or PTH1-34 (purple transparent ribbon, PDB 8FLQ), and Gs (cyan and blue, respectively). PCO371 is shown as an orange stick representation within the TMD, while PTH1-34 is shown as a yellow ribbon in the orthosteric site. The main PTH1R structural differences are highlighted in red dashed rectangles; in complex with PCO371, the TM6 sharp kink assumes a linear short linker conformation, the extracellular end of TM6 is fully helical, and the TM1/TM7 domain is less bent, compared to PTH1R in complex with PTH1-34. **B-D)** PCO371’s possible tautomeric forms at the 4-dihydro-imidazolone group. **E)** Structural details of PCO371 interactions between TM6 and PCO371’s 1,4-dihydro-imidazolone group, as reported in PDB files 8JR9 and 8GW8; PDB resolution is reported. **F)** occupancy of hydrogen bonds between the TM6 backbone and PCO371; **G)** RMSD analysis of PCO371 keto A and enol forms, bound to PTH1R, during MD simulations in the absence of Gs; **H**) TM6-PCO371 distance analysis during MD simulations.

#### 3.4.2. Assessing PCO371 tautomeric isomers when bound to PTH1R

The structure of PCO371 is characterised by a 1,3,8-triazaspiro[4.5]dec-1-en-4-onespirocycle linked to a substituted benzene ring via an ethylsulfonyl linker, with hydantoin and 4-(trifluoromethoxy)phenyl terminal groups (Figure 7A). In complex with PTH1R and Gαs (PDBs 8JR9 and 8GW8), PCO371 engages and stabilises TM6 primarily via its 1,4-dihydro-imidazolone group (Figure 7B). The chemistry of this heterocyclic ring allows three different tautomeric forms (Figure 7B-D). The keto tautomer A (Figure 7C) is the most stable and largely prevalent form in aqueous solution (37); the enol (Figure 7D) and keto tautomer B (Figure 7B) possess much lower stability. The keto tautomer B is the least stable due to fewer conjugation effects. However, the stability of PCO371 tautomers can be influenced by the local protein environment provided by PTH1R, such that the form responsible for binding and agonism is not necessarily the most stable in aqueous solution. The analysis of the donor-acceptor distances (Figure 7E) between the 1,4-dihydro-imidazolone and TM6, as reported in the PDB files 8JR9 and 8GW8, suggests weak hydrogen bonds with the backbone nitrogen atoms of L416, F417 and PCO371 (distances between 3.12 A and 3.60 A), whereas strong hydrogen bonds are usually < ∼3.2 A (38). Such relaxed donor-acceptor distances and the simultaneous presence of two hydrogen-bond donors on the PTH1R backbone (i.e. L416 and F417 nitrogen atoms) do not rule out PCO371 engaging PTH1R as a mix of keto A and enol forms.

We performed molecular dynamics (MD) simulations of PTH1R in complex with keto A and enol PCO371 forms, in the absence of Gαs, to retrieve structural insights into the most stable tautomer bound to the receptor. Both tautomers displayed similar root-mean-square deviation (RMSD; Figure 7G) and a comparable tendency to dissociate from TM6 (computed as the distance between the L416 Cα atom and PCO371’s 1,4-dihydro-imidazolone group, Figure 7H). However, the enol tautomer did not form hydrogen bonds with either L416 or F417 backbone nitrogen atoms, while keto A PCO371 engaged F417 in a low-occupancy interaction (∼15% of MD frames, Figure 7F). In light of this slightly higher consistency with the experimental structures, we considered PCO371 in keto form A for the MD simulations successively performed.

#### 3.4.3. PCO371 requires G-protein engagement to form a stable complex

To test whether PCO371 can stabilise an active receptor conformation independently, we performed MD simulations of PTH1R:PCO371 with and without any G-protein (Figure 8A), PTH1R:PCO371:Gαs (Figure 8B), PTH1R:PCO371:Gαi3 (Figure 8C), and PTH1R:PCO371:Gαq (Figure 8D). During three independent 1 µs MD replicas, PCO371 in the absence of a G-protein was unstable and completely dissociated in two of three replicas (Figure 8A). This suggests that PCO371 binds to a preassembled but not yet functional PTH1R:Gαs complex, rather than forming a stable complex with the receptor ahead of Gαs engagement.

**Figure 8.**
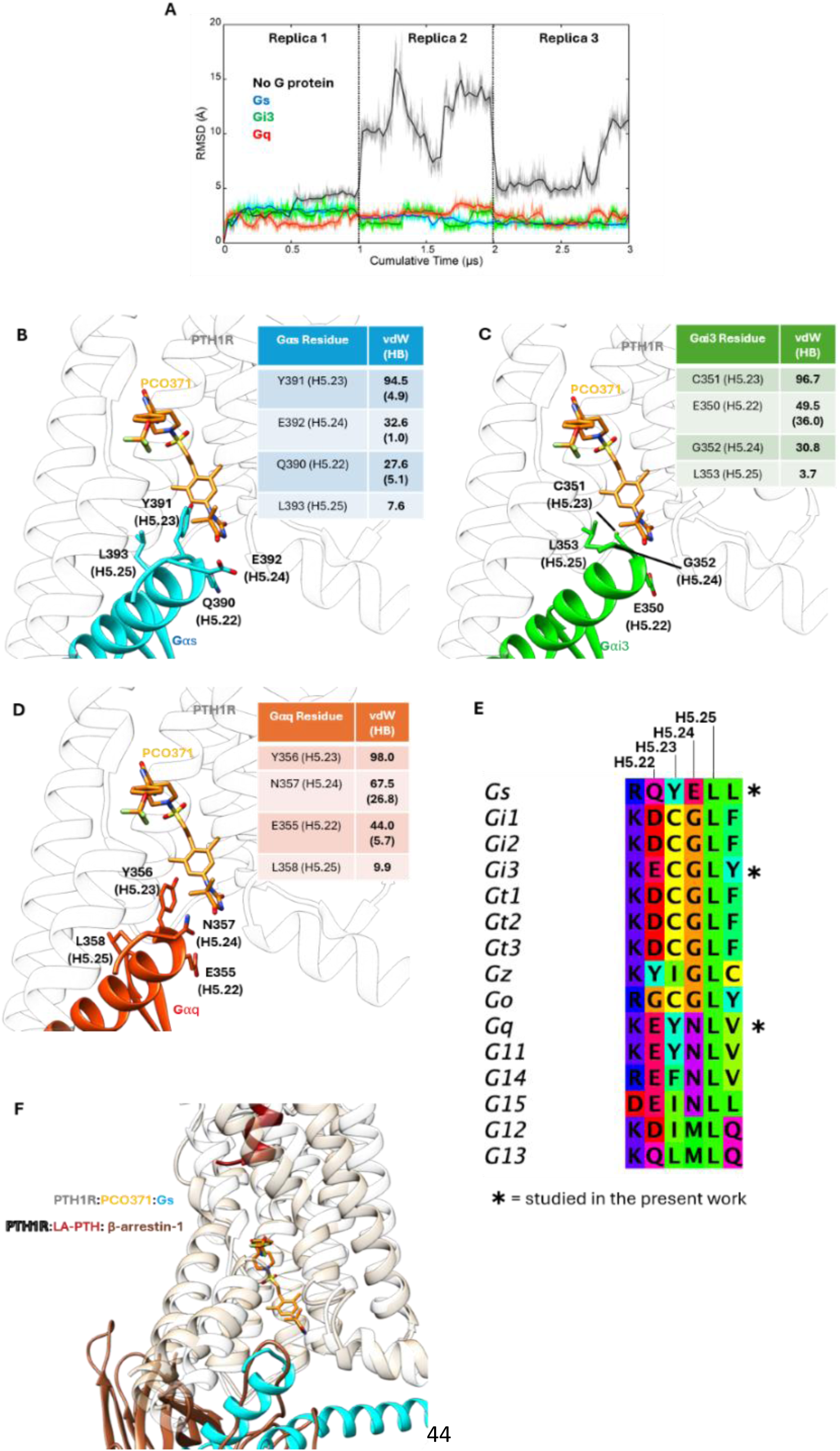
Structural basis of PCO371-mediated PTH1R activation. **A)** RMSD of PCO371 during MD simulations in complex with PTH1R and Gs (blue line, three replicas), Gi3 (green, three replicas), Gq (red, three replicas), or without G protein (black). (**B– D)** Representative configurations of PCO371 in complex with Gs (cyan ribbon), Gi3 (green), Gq (red); G proteins’ residues involved in interactions with PCO371 are reported in the inserted tables (van der Waals (vdW) and hydrogen bonds (HB) are reported as a percentage of MD frames) and shown in stick. **E**) Multiple sequence alignment of the C-terminal α5-helix region (positions H.5.22-25) across Gα-protein families. Residues are coloured according to amino acid identity. **F**) Comparison between PTH1R in complex with PCO371 (white ribbon, PDB 8JR9) or long-acting (LA)-PTH1-34 (tan transparent ribbon), and Gs (cyan ribbon) of β-arrestin-1 (maroon ribbon, PDB 9LZ0). β-arrestin-1 finger loop backbone overlaps the Gs helix 5. PCO371 2D structure is shown in Figure 7A.

#### 3.4.4. G-protein α5 helix residues contribute to subtype potency differences for PCO371

In all G-protein complexes: PTH1R:PCO371:Gαs, PTH1R:PCO371:Gαi3 and PTH1R:PCO371:Gαq, PCO371 remained stably bound to both PTH1R and the associated G-protein (Figure 8A). In all complexes, PCO371 formed extensive interactions with PTH1R (Tables 4 and 5). Four residues from the C-terminal α5 helix (H5) of the G-protein, H5.22, H5.23, H5.24, and H5.25 (G-protein numbering system), contributed to PCO371 stabilisation (Figure 8B-D). Among these, H5.23 showed the highest contact frequency (Y391 in Gαs, Y356 in Gαq, and C351 in Gαi3), followed by H5.24 (E392, N357, and G352, respectively) and H5.22 (Q390, E355, and E350). Gαs engaged PCO371 in fewer contacts and hydrogen bonds (Table 4, Figure 7A) than Gαq and Gαi3. Gαq formed more stable interactions with H.24 (N357) and H.22 (E355) than Gαs and Gαi3. PCO371 formed a hydrogen bond with the Gαi3 backbone at E350 (∼36% of MD frames).

**Table 4.** PTH1R residues involved in van der Waals interactions with PCO371 in complex with Gαs-, Gαq-, and Gαi3-proteins. Occupancies are expressed as % of MD frames. Residues are coloured according to their structural location within the receptor.

| PTH1R | Gas | Gaq | Gai3 |
| --- | --- | --- | --- |
| TYR459(7.57) | 100 | 99.1 | 99.4 |
| PHE417(6.49) | 97.5 | 96.3 | 99.6 |
| ARG219(2.46) | 95.2 | 36.5 | 37.2 |
| GLU302(3.50) | 90.9 | 79 | 90.6 |
| VAL412(6.44) | 85.6 | 93.7 | 91 |
| PHE454(7.52) | 84.3 | 72.8 | 77.9 |
| HIS223(2.50) | 80.7 | 67.9 | 88.3 |
| ILE458(7.56) | 77.7 | 83.6 | 71.1 |
| MET414(6.46) | 68.5 | 96.5 | 95.1 |
| LEU416(6.48) | 66.1 | 95.5 | 95.8 |
| LEU413(6.45) | 65.1 | 82.8 | 63.8 |
| ASN463(8.47) | 53.5 | 65.5 | 77.4 |
| GLN451(7.49) | 43.6 | 16.9 | 32.9 |
| PRO415(6.47) | 40.5 | 55.3 | 47.5 |
| ILE299(3.47) | 39.5 | 18.2 | 41.2 |
| PHE447(7.45) | 34.9 | 12.8 | 24.9 |
| GLY464(8.48) | 26.8 | 29.5 | 1.7 |
| VAL455(7.53) | 22.6 | 7.4 | 12.9 |
| LEU226(2.53) | 17.5 | 14 | 43.5 |
| PHE230(2.57) | 17 | 7.8 | 0 |
| LEU306(3.54) | 6.9 | 24.5 | 10.5 |
| CYS462(7.60) | 6.8 | 12.2 | 1.5 |
| LEU411(6.43) | 5.3 | 20.2 | 19.6 |

**Table 4.** PTH1R residues involved in van der Waals interactions with PCO371 in complex with Gas-, Gaq-, and Gai3-proteins. Occupancies are expressed as % of MD frames. Residues are coloured according to their structural location within the receptor.
| Colour Key |
| --- |
| ECD |
| TM1 |
| TM2 |
| TM3 |
| TM4 |
| TM5 |
| TM6 |
| TM7 |
| Helix-8 |
| ICL1 |
| ICL2 |
| ICL3 |
| ECL1 |
| ECL2 |
| ECL3 |

**Table 5.** PTH1R residues involved in hydrogen-bond (side-chain or backbone) interactions with PCO371 in complex with Gαs-, Gαq-, and Gαi3-proteins. Occupancies are expressed as % of MD frames. Residues are coloured according to their structural location within the receptor.

| PTH1R | Gas | Gaq | Gai3 |
| --- | --- | --- | --- |
| ARG219(2.46) | 94.4 | 35.2 | 17.1 |
| PHE417(6.49) | 68 | 90.2 | 96.4 |
| TYR459(7.57) | 59.9 | 79.8 | 61.7 |
| GLN451(7.49) | 28.3 | 5.1 | 9.3 |
| GLY464(8.48) | 25.5 | 28.3 | 1.6 |
| LEU416(6.48) | 3.7 | 12.2 | 10.3 |
| ASN463(8.47) | 2.9 | 5.8 | 19.8 |

Despite some variability inherently due to the different sequences across the different G-proteins, PCO371 formed stable interactions with both PTH1R and the Gα subunits C-terminal, consistent with our functional and TRUPATH assays. The sequence analysis of H5 across all the G-proteins suggests compatibility with a broader range of Gα subtypes (Figure 8E), especially Gαi1, Gαi2, Gα11, and Gα14, due to the high sequence similarity at positions H.5.22-25, relative to Gαs, Gαq, and Gαi3.

#### 3.4.5. Structural validation of β-arrestin engagement by PCO371

Superposition of the PTH1R:PCO371:Gαs structure (PDB 8JR9) with the recently determined PTH1R:LA-PTH1:β-arrestin-1 structure (PDB 9LZ0) with β-arrestin-1 in ‘core’ engagement (39) reveals no significant steric clashes between β-arrestin and PCO371 (Figure 8F). Notably, the finger loop of β-arrestin, which inserts into the receptor core and engages the region surrounding the PCO371 binding site, is conserved between β-arrestin-1 and -2.

This suggests that PCO371 binding is compatible with β-arrestin recruitment and supports the observed activation of β-arrestin-2 by PCO371 (Figure 6).

## 4. DISCUSSION

This study presents a detailed investigation into the molecular mechanisms of activation of the PTH1R by two ligands with very different activation strategies; PCO371, a small molecule non-peptide that binds to the intracellular face of the receptor and PTH 1-34, a peptide ligand that binds to the extracellular orthosteric site. PCO371 was previously reported to be incapable of stabilising an active PTH1R conformation that couples to β-arrestin-2 (12,13). The current study establishes that PCO371 can recruit β-arrestin-2, albeit with lower E_max_ than PTH 1-34 and only under high receptor reserve. This dependence on receptor density reflects low intrinsic efficacy rather than signalling bias. A truly biased ligand would fail to activate a pathway regardless of receptor reserve, as seen for orforglipron at the GLP1R (9). In contrast, PCO371 behaves as a balanced partial agonist across all signalling pathways examined at the PTH1R.

Functionally, PCO371 was unable to stabilise the high-affinity, mini-Gαs–coupled conformation of membrane-bound PTH1R, in contrast to PTH 1-34. Its functional affinity derived from both Operational Model and Furchgott analyses, converged on a dissociation constant that matched its pEC50 under low-receptor-reserve conditions, a defining feature of partial agonism. Receptor-reserve analysis (Figure 2) further demonstrated a linear occupancy-response relationship for PCO371 in cAMP accumulation, whereas PTH 1-34 retained high efficacy despite receptor depletion. These findings show that the signalling output of PCO371 is strongly dependent on receptor abundance and coupling efficiency.

Rather than a limitation, this possibly reflects the mechanistic signature of intracellular agonism: PCO371 stabilises the active ternary complex only when coupling conditions are optimal. Physiologically, tissues with high PTH1R expression, such as bone and kidney (40), are therefore more likely to support PCO371 responses, whereas low-expression environments may constrain its potency.

Interestingly, the differences in raw BRET signal between PTH 1-34 and PCO371 were evident in TRUPATH assays (Figure 4), potentially showing how the kinetics of GαsL-protein activation differ between the two ligands. The slower activation kinetics observed for PCO371, compared with the rapid G-protein subunits dissociation induced by PTH 1-34, provide important mechanistic insight into how the small molecule engages the receptor. PTH 1-34 activates PTH1R through the canonical extracellular route, producing an immediate outward movement of TM6 and rapid G-protein engagement. In contrast, PCO371 must first access its intracellular binding pocket and can only form a stable complex once the receptor is already pre-coupled to a G-protein, as observed in our MD simulations in which PCO371 dissociates from uncoupled receptor states (Figure 8A). Structurally, PCO371 acts as a “molecular wedge” that inserts into the intracellular groove of PTH1R (12); its engagement stabilises a distinct active-state geometry characterised by a straighter TM6 mid-segment and a more compact TM1–TM7 bundle. This structural arrangement points towards a mechanism in which PCO371 reinforces, rather than initiates, the receptor-G-protein interface, explaining the kinetic lag before steady-state signalling is reached. Possibly PCO371 behaves analogously to cytoplasmic lipids that stabilise preassembled complexes (41), but with the unique capacity to act as an agonist.

Pharmacological antagonism further supports distinct binding modes between the peptide and small-molecule agonists. DS69910557 behaved as a high-affinity antagonist for peptide-mediated signalling, also reported previously (35), but DS69910557 did not inhibit PCO371-stimulated cAMP accumulation, confirming that the two agonists occupy separate receptor binding sites. The absence of antagonism toward PCO371-mediated signalling, together with the lack of structural data describing how DS69910557 exerts its inhibitory effect, suggests that DS69910557 primarily engages the receptor’s N-terminal extracellular domain rather than the transmembrane bundle. This interpretation is consistent with the intracellular binding mode for PCO371, which acts through a distinct allosteric site to stabilise receptor-G-protein coupling and drive activation independently of extracellular antagonists.

Differences in signalling can also be observed in how PTH1-34 activates different G-protein subtypes. For GαsL-protein, PTH 1-34 produces a large, rapid decrease in BRET, consistent with efficient heterotrimer dissociation and potentially reflecting a degree of pre-coupling between receptor and G-protein (42,43), though this does not explain the lag-phase seen for PCO371. For Gαi3-protein, heterotrimer dissociation is of consistent magnitude across all time points. This suggests a modulatory role for Gαi3 at the PTH1R rather than a major driver of signalling; importantly, coupling of PTH1R to Gαi has been reported previously (44,45). Activation of Gαq(R183Q)-protein gives a third pattern, with activation increasing over time to a stable maximum. These profiles show that PC0371 activates all tested G-protein subtypes, arguing against meaningful G-protein bias.

Structural modelling and MD simulations support a scenario in which PCO371 requires the G-protein for complete stabilisation within the PTH1R intercellular binding site and to stabilise the open conformation of TM6. Once a ternary complex with PTH1R and Gα-protein is formed, PCO371 is consistently stable within the chemical environments provided by different Gα subunit C-termini.

Collectively, our data demonstrate that PCO371 requires preassembled G-protein engagement and high receptor reserve to stabilise a distinct active-state geometry of PTH1R. This conformation supports activation of Gαs, Gαi3, Gαq, and β-arrestin pathways but yields globally reduced signalling efficiency compared with PTH 1-34. Importantly, PCO371’s intracellular binding mode renders its activity insensitive to antagonism by extracellular R₀-targeting compounds, confirming that it operates through a mechanistically independent route. These findings provide crucial mechanistic insight into an atypical yet therapeutically valuable small-molecule modulator of class B1 GPCR activity, establishing a functional foundation for the rational design of next-generation targeted therapeutics.

## Acknowledgements

Work in G.L.’s lab is funded by the BBSRC (BB/Y513817/1, BB/W014831/1, and UKRI1925.). We also thank the David James Trust for funding. G.L. was a Royal Society Industry Fellow (INF/R2/212001). G.D. acknowledge funding from Leverhulme Trust (RL-2025-012).

## CRediT authorship contribution statement

Farhaan Napier-Khwaja: Writing - review & editing, Validation, Methodology, Investigation, Formal analysis, Conceptualisation.

Zamara Mariam: Validation, Methodology, Investigation, Formal analysis.

Yasaman Abdolhay: Validation, Methodology, Investigation, Formal analysis.

Theo Redfern-Nichols: Writing - review & editing, Validation, Methodology, Investigation, Formal analysis.

Graham Ladds: Writing - review & editing, Validation, Methodology, Resources, Supervision.

David R. Poyner: Writing - review & editing, Validation, Methodology, Supervision, Conceptualisation.

Giuseppe Deganutti: Writing - review & editing, Validation, Methodology, Formal analysis, Supervision, Resources, Funding acquisition, Conceptualisation.

Mark Wheatley: Writing - review & editing, Validation, Methodology, Formal Analysis, Supervision, Resources, Funding acquisition, Conceptualisation.

Hoor Ayub: Writing - Original draft, Validation, Methodology, Formal Analysis, Supervision, Resources, Funding acquisition, Conceptualisation.

## Declaration of competing interest

The authors declare no competing financial interest.

## Data Availability statement

All data reported in this paper will be shared by the corresponding author upon reasonable request. MD simulation trajectories have been deposited in the public repository Zenodo at https://zenodo.org/records/226580.

## Abbreviations

ANOVA: analysis of variance
AUC: area under the curve
BODIPY: boron-dipyrromethene fluorophore
BRET: bioluminescence resonance energy transfer
cAMP: cyclic adenosine 3’, 5’ monophosphate
ECD: extracellular domain
ECL: extracellular loop
GnTI^-^: N-acetylglucosaminyltransferase I deficient
GPCR: G-protein-coupled receptor
G-protein: guanine nucleotide binding protein
HEK 293: human embryonic kidney cell line
HTRF: homogenous time-resolved fluorescence
IBMX: 3-isobutyl-1-methylxanthine
MD: Molecular dynamics
Mini-G_α_: engineered GTPase domain of G_α_ subunit
NanoBRET: NanoLuciferase Bioluminescence Resonance Energy Transfer
NLuc: nanoluciferase
PTH: parathyroid hormone
PTH1R: parathyroid hormone 1 receptor
RMSD: root-mean-square deviation
SEM: standard error of the mean
tetO: tetracycline-operator sequence
tetR: tetracycline-repressor
TR-FRET: time-resolved Förster resonance energy transfer

## Notes

### Competing Interest Statement

The authors have declared no competing interest.

### Summary of Updates

Updated the figures and added co-authors (who contributed with additional data).

